# A machine-learning algorithm correctly classifies cortical evoked potentials from both natural retinal stimulation and electrical stimulation of the optic nerve

**DOI:** 10.1101/2021.03.27.437330

**Authors:** Vivien Gaillet, Elodie Geneviève Zollinger, Diego Ghezzi

**Affiliations:** Medtronic Chair in Neuroengineering, Center for Neuroprosthetics and Institute of Bioengineering, School of Engineering, École polytechnique fédérale de Lausanne, Switzerland

## Abstract

**Objective:** Optic nerve’s intraneural stimulation is an emerging neuroprosthetic approach to provide artificial vision to totally blind patients. An open question is the possibility to evoke individual non-overlapping phosphenes via selective intraneural optic nerve stimulation. To begin answering this question, first, we aim at showing in preclinical experiments with animals that each intraneural electrode could evoke a distinguishable activity pattern in the primary visual cortex.

**Approach:** We performed both patterned visual stimulation and patterned electrical stimulation in healthy rabbits while recording evoked cortical activity with an electrocorticogram array in the primary visual cortex. Electrical stimulation was delivered to the optic nerve with the intraneural array OpticSELINE. We used a support vector machine algorithm paired to a linear regression model to classify cortical responses originating from visual stimuli located in different portions of the visual field and electrical stimuli from the different electrodes of the OpticSELINE.

**Main results:** Cortical activity induced by visual and electrical stimulation could be classified with nearly 100% accuracy relative to the specific location in the visual field or electrode in the array from which it originated. For visual stimulation, the accuracy increased with the separation of the stimuli and reached 100% for separation higher than 7 degrees. For electrical stimulation, at low current amplitudes, the accuracy increased with the distance between electrodes, while at higher current amplitudes, the accuracy was nearly 100% already for the shortest separation.

**Significance:** Optic nerve’s intraneural stimulation with the OpticSELINE induced discernible cortical activity patterns. These results represent a leap forward for intraneural optic nerve stimulation towards artificial vision.

## Introduction

Blindness is a medical condition impairing the quality of life of the affected people and their relatives; hence it represents a high medical, economic and social cost for society. While therapies are available for several pathologies causing vision loss and blindness, some others currently have no treatment [1]. Among these diseases, retinitis pigmentosa is a set of inherited retinal rod-cone dystrophies with a prevalence of about 1:4,000 individuals causing the progressive loss of rod photoreceptors, loss of night vision and the constriction of the visual field (tunnel vision), later followed by cone dysfunctions and eventually profound blindness [2]. The degeneration of retinal photoreceptors characterises retinitis pigmentosa. Consequently, the other retinal neurons (i.e. bipolar cells, amacrine cells and retinal ganglion cells) are generally considered to be preserved and to form a functional network that can be activated via electrical stimulation evoking the sensation of discrete points of light called phosphenes. Artificial vision is the perception of the world using an ensemble of these phosphenes. Retinal prostheses have been extensively investigated in animal experiments [3–9] and in retinitis pigmentosa patients to fight blindness [10–12]. However, none of these devices is available in clinics today, and the few that were available have been discontinued.

Large clinical studies with retinal implants in retinitis pigmentosa patients showed a large performance variability [13–15]. The number of functional electrodes able to elicit phosphenes with the Argus® II retinal implant was variable, as low as 5 out of 60 [13]. Postoperative outcomes were also variable, from just phosphenes perception to object recognition and reading capability [13]. Similar variable outcomes were observed in retinitis pigmentosa patients implanted with the Alpha-IMS/AMS subretinal prostheses, ranging from just light perception to restoration of partial visual acuity [15]. Variable perceptual outcomes could be attributed to the advance stage of retinal degeneration. In late-stage retinitis pigmentosa patients, post-mortem retina analysis revealed that 78% of bipolar cells and 30% of retinal ganglion cells are spared, while it was the case for only 5% of photoreceptors [16–19]. However, there are divergent opinions about the functional integrity of retinal circuits. While some studies reported the anatomical and functional preservation of retinal ganglion cells in both humans and animal models [16,20,21], some others documented the remodelling and reorganisation of the synapses in the inner nuclear layer [22,23], which might compromise the efficacy of electrical stimulation [24–27]. The deafferentation of bipolar cells caused by photoreceptors’ death is followed by a negative plasticity period, which is not fully understood. Changes in synaptic connectivity might form aggregates of synaptic connexions between survival cells, known as microneuromas, which are presumably the source of the spontaneous retinal hyperactivity [24,26,28]. These plastic events might create connectivity loops, reducing the retinal excitability and impairing the ability to encode visual information with retinal implants [29]. Retinal network-mediated stimulation in retinitis pigmentosa patients is also subjected to rapid phosphene’s fading due to static adaptation and bipolar cells’ desensitisation [30,31]: a phenomenon requiring cognitively exhausting large head and body movements to counteract it. Therefore, bypassing the retinal network might be an attractive strategy to reduce the variability in retinal stimulation. One approach was developed by the AV-DONE project based on an intrapapillary electrode array [32,33]. A retinitis pigmentosa patient implanted with three wires reported the perception of distinguishable phosphenes upon stimulation of the optic disc [34].

Long-term mechanical stability of epiretinal prostheses is another critical element leading to an increase in the stimulation threshold associated with the increase of the electrode-to-retina distance [35,36]. Also, several exclusion criteria limit the eligibility for retinal prostheses, as appeared from enrolment criteria in several clinical trials, and here summarised: retinal detachment or trauma, sub-macular choroidal neovascularization, extremely thin conjunctiva, too small or too long axial length, corneal ulcers, abnormalities in the typical curvature of the retina, eyeball trauma, and corneal opacity or lack of optical transparency preventing adequate visualisation of the inner structures of the eye or impeding the correct placement of the device. Retinal prostheses might not be limited by all those criteria simultaneously since some of them might apply only to specific implants. However, optic nerve’s direct electrical stimulation appears an attractive strategy to circumvent all the above-mentioned limitations in a single step. Another consideration is the restored visual angle. Previous studies pointed out that retinitis pigmentosa patients will benefit from the restoration of a wide visual angle [37,38]. Although some wide-field retinal prostheses are under development [5,39–42], optic nerve’s stimulation is attractive because its relatively small diameter facilitates the electrical stimulation of a wide visual field.

Early studies on optic nerve’s stimulation were performed in retinitis pigmentosa patients [43,44]. The first subject received a 4-contacts self-sizing cuff electrode around the intracranial segment of the optic nerve. Following psychophysical testing, the subject correctly identified basic shapes and character-like symbols, even if the phosphenes were reported as unstable, possibly due to the device’s movement caused by the poor anchorage of the cuff electrode. Following these results, the C-sight project proposed an optic nerve prosthesis based on a few (3 to 5) penetrating rigid and stiff microelectrodes and tested the concept in preclinical experiments [45,46]. An optic nerve prosthesis (OpticSELINE) was also proposed by our group [47]. Compared to other approaches, the OpticSELINE provides higher mechanical stability due to its three-dimensional anchoring wings, better mechanical compliance because of thin-film microfabrication on polymeric materials and higher electrodes’ number because of the high integration capacity of cleanroom processes. Despite early clinical studies in humans [34,43,44] and preclinical experiments in animals [45–47], optic nerve stimulation is not mature yet for clinical use. Several technical improvements and validations are still required, such as electrode miniaturisation, increase of the electrodes’ number and density, long-term biocompatibility assessment in the optic nerve and demonstration of localised and non-overlapping phosphenes when multiple electrodes are activated simultaneously or in rapid succession. A step towards this latter goal is to show that the stimulation through different electrodes induces distinguishable cortical activity patterns, suggesting that different fibres of the optic nerve are recruited, and non-overlapping phosphenes might be evoked.

Preclinical studies in animals often rely on the peak-to-peak analysis of electrocorticogram (ECoG) recordings to determine the portion of the visual cortex activated by the stimulation. However, ECoG recordings with microelectrodes are often limited by low spatial resolution. A recent report found that the cortical tissue around an ECoG electrode that contributes to its activity is approximately 3-mm in diameter [48], most likely because of the volume conduction of the potentials. The mapping of spatially organised cortical activity patterns in relatively small animals could be hindered, if not prohibited, by this physical limitation. On the other hand, a previous study presenting letters to a rhesus monkey demonstrated that machine-learning algorithms might overcome this problem since they benefit from the information of the whole spectrogram of ECoG recordings [49]. Therefore, we propose a machine-learning approach not relying exclusively on the peak-to-peak amplitude but encompassing the whole signal spectrogram to correctly classify and predict the ECoG signal originating from patterned visual stimulation and optic nerve’s intraneural electrical stimulation. First, we validated the algorithms using patterned visual stimuli. Then, we demonstrated that the cortical signals evoked by electrical stimulation from different electrodes of the OpticSELINE were correctly classified and predicted.

These results represent a leap forward for intraneural optic nerve stimulation towards an artificial vision. Also, this approach might have wide use in neuroprosthetics for the preclinical validation of visual prostheses and other sensory prostheses.

## Materials and methods

### Animal handling and surgery

Animal experiments were performed according to the authorisation GE519 approved by the Département de l'Emploi, des Affaires Sociales et de la Santé, Direction Générale de la Santé of the Republique et Canton de Genève (Switzerland). Female rabbits (>16 weeks, >2.5 kg) were premedicated 30 min before the transfer to the surgical room with an intramuscular injection of xylazine (3 mg kg^−1^; Rompun® 2%, 20 mg ml^−1^, 0.15 ml kg^−1^), ketamine (25 mg kg^−1^; Ketanarkon® 100 mg ml^−1^, 0.25 ml kg^−1^) and buprenorphine (0.03 mg kg^−1^; Temgesic® 0.3 mg ml^−1^, 0.1 ml kg^−1^). Rabbits were left under 100% O_2_. 10 to 15 min after premedication, a 22G catheter was placed in the ear’s marginal vein. Local anaesthesia was provided to the throat (Xylocaine 10%, spray push). The rabbit was intubated with a 3.5-mm tracheal tube with a balloon and ventilated (7 ml kg^−1^, rate: 40/min, positive end-expiratory pressure: 3 cm of H_2_0). Eye gel (Lacrigel) was placed on the eye to protect it from drying. Anaesthesia and analgesia were provided intravenously with propofol (10 mg kg-1 h^−1^; 20 mg ml^−1^, 0.5 ml kg^−1^ h^−1^) and fentanyl (0.005 mg kg−1 h^−1^; 0.05 mg ml^−1^, 0.1 ml kg^−1^ h^−1^). Rabbits were placed on a heating pad at 35 °C for the entire surgery. The depth of anaesthesia was monitored continuously throughout the procedure, including heart rate, temperature and pulse oximetry. Fluid supplementation was administered intravenously to prevent dehydration (physiological solution 20-35 ml kg^−^ 1, IV). The rabbit’s head was shaved and secured gently within a stereotactic frame (David Kopf Instruments). Before cortical skin incision, lidocaine (6 mg kg^−1^; Lidocaine 2%, 20 mg ml^−1^, 0.3 ml kg^−1^) was injected subcutaneously at the surgical sites. After 5 min, the skin was opened and pulled aside to clean the skull with cotton swabs. A craniotomy was made to expose the visual cortex. A 32-channel epidural ECoG array (E32- 1000-30-200; NeuroNexus) was placed over V1. For electrical stimulation, a frontotemporal craniotomy was made to access the optic nerve. The intraneural electrode array was inserted in the optic nerve in the pre-chiasmatic area. The surgical implantation was performed by piercing the nerve with a needle (nylon black DLZ 4,8-150 10/0, FSSB) and by guiding the intraneural electrode array transversally into the nerve. All rabbits were euthanised at the end of the acute recording procedures while still under anaesthesia, with an intravenous injection of pentobarbital (120 mg kg^−1^).

### Electrophysiological recording and stimulation

For patterned visual stimulation, white rectangles were projected (ML750ST; Optoma) on a reflective screen placed approximately 90 cm away from the rabbit. The projected area covered 28 x 44 degrees of the visual field, and the full luminance of the projector was 530 cd m^−2^. Rectangles of 1/4^th^ of the screen’s size (14 x 22 degrees) were projected in 10 different locations along the horizontal meridian separated by 2.2 degrees. Each stimulus lasted for 16 ms (based on the projector frame rate). Sixty flashes were presented for each stimulus location. For patterned electrical stimulation, two microelectrode arrays were used: Flat-OpticSELINE and OpticSELINE. The OpticSELINE was previously described by our group [47]. Briefly, it is a polyimide-based looped microelectrode array. Six microelectrodes (0.0078-mm^2^ area) are located on each side of the OpticSELINE, with a reference electrode and a ground electrode placed outside the active area. Two three-dimensional flaps extend from both sides of the main body and carry two electrodes each; two more electrodes are located on each side of the main body. The dataset with the OpticSELINE was obtained from the previous experiment [47], during which only the six electrodes on the top side were used for stimulation. The Flat-OpticSELINE is a modified version of the OpticSELINE. The array is a single polyimide strip with a linear and planar distribution of eight electrodes (40-μm diameter, 160- μm pitch, 0.0013-mm^2^ area) with a reference electrode and a ground electrode placed outside the active area. The microelectrode array was attached to a current stimulator (IZ2MH; Tucker-Davis Technologies). Optic-nerve stimulation was performed with 150-μs biphasic cathodic-first current pulses (asymmetric: 750-μs anodic phase at one-fifth of the cathodic amplitude) at various cathodic current amplitudes (10, 25, 50, 75, 100, 150, 200, 250, 500, 750, 1,000, 1,500 and 2,000 μA) delivered in a scrambled manner. For each electrode and current amplitude, 30 stimuli were delivered with the Flat-OpticSELINE and 10 with the OpticSELINE. For cortical recordings, an ECoG array composed of 32 (4 × 8) platinum electrodes with a diameter of 200 μm and a pitch of 1 mm (E32-1000-30-200; NeuroNexus) was placed over V1 contralateral to the stimulated eye and connected to an amplifier (PZ5; Tucker-Davis Technologies) via a 32-channel analogue head stage (ZIF-Clip Analog Headstage; Tucker-Davis Technologies). Data were filtered between 0.1 and 500 Hz and digitised at 12 kHz. Epochs synchronous to the stimulation’s onset (from –100 ms to +750 ms) were then extracted from the data stream and analysed with MATLAB (MathWorks).

### Signal processing and feature extraction

The ECoG signal was band-pass filtered between 3 and 500 Hz, then offset by a value equal to the average voltage of the 100-ms preceding the stimulus’s onset. The signal spectrogram was computed with a built-in MATLAB function (continuous 1-D wavelet transform) using the Morlet wavelet to extract the frequency bands from 10 to 300 Hz. The wavelet power was obtained by squaring two times the magnitude of the wavelet transform. The wavelet power was then normalised by subtracting the average power of the 100 ms before the stimulus’s onset and then dividing the result by the same value for each frequency bands (baseline-correction). Nine classes of features were extracted from the time course portion from 5 ms after the stimulus’s onset (to remove the stimulus artefact that occurs between 0 to 4 ms) to 100 ms after the stimulus. The continuous features named ‘Time’ and ‘Slope’ consist of the time course itself (‘Time‘) and its derivative (‘Slope’). The punctual features named ‘PA’ and ‘PL’ consisted of the VEP signal’s peak amplitude (‘PA‘) and the positive peak’s latency (‘PL‘). The broad-band gamma power (‘BGP‘) was obtained by averaging the frequency bands from the wavelet power ranging from 40 to 150 Hz. The punctual features in the frequency domain named ‘Power Amplitude’, ‘Power Frequency’ and ‘Power Timing’ correspond respectively to the maximum value of the power wavelet spectrogram, its frequency and its timing. Finally, the wavelet power spectrogram was down-sampled in the time axis from 12 kHz down to 1200 Hz and used as the continuous feature named ‘Power’.

### Classification

For patterned visual stimulation and patterned electrical stimulation with the Flat-OpticSELINE, both a classification and a linear regression were performed. For patterned electrical stimulation with the OpticSELINE, only the classification was performed. The classification was performed in MATLAB using a support vector machine algorithm (SVM), determining the hyperplane in the feature space that maximises the margins (the projection of the data points on the hyperplane). The SVM algorithm was used with the MATLAB error-correction output codes allowing the SVM classifier to be used on a multi-class data set. No kernel function was used. We ran 6-fold cross-validation, where 5/6 of the observations were used for training (training set) and 1/6 for testing (testing set). The classification was repeated on 6 different pairs of training and testing sets. The final classification accuracy score was the average of the 6 trials. Features’ selection was performed on the training set only. We first conducted the classification on all the stimuli at once for patterned visual stimulation and patterned electrical stimulation with the Flat-OpticSELINE. Then, we computed the classification accuracy between pairs to evaluate the impact of the distance between stimuli or electrodes on the classification accuracy. For patterned electrical stimulation with the OpticSELINE, the classification was conducted on all the stimuli at once only.

### Linear regression

The correlation coefficient between the features and the stimuli’ location was computed, and a regression model was built with the features with the highest absolute correlation coefficient value. The model takes the form of y = β_0_+β_1_X_1_+β_2_X_2_+β_3_X_3_+ … + β_n_X_n_ + ɛ, where X_1_ to X_n_ are the features used as input, β_0_ to β_n_ are the weights of the features, ɛ is the error term, and y is the predicted location of the rectangle’s centre or the electrode’s position. The observations were split into training and testing sets of a size equal to 5/6 and 1/6 of the total number of observations, which was repeated 60 times for patterned visual stimulation and 30 times for patterned electrical stimulation with the Flat-OpticSELINE to gradually shift the testing sets by an increment equal to the number of stimulations. The average model predictions and the root-mean-square error (RMSE) were reported as the average values of the different splits.

## Results

### Classification of visual responses

We took advantage of patterned visual stimulation to validate the SVM classification algorithm on ECoG recordings (**Figure 1**). An anaesthetised rabbit was placed in front of a screen (**Figure 1a**), and white flash stimuli (14 x 22 degrees rectangles) were projected on dark background in 10 different locations along the horizontal meridian (red dots in **Figure 1b**). Visually evoked potentials (VEPs) elicited by patterned visual stimulation (**Figure 1c**) were detected using the ECoG electrode array placed over V1 contralateral to the stimulated eye. Strong patterned flash stimulation was selected among the various stimuli used in vision research to mimic the strong and fast activation induced by electrical stimulation as closely as possible. Flashes and electric pulses are both fast transients, not physiological in normal vision.

**Figure 1.**
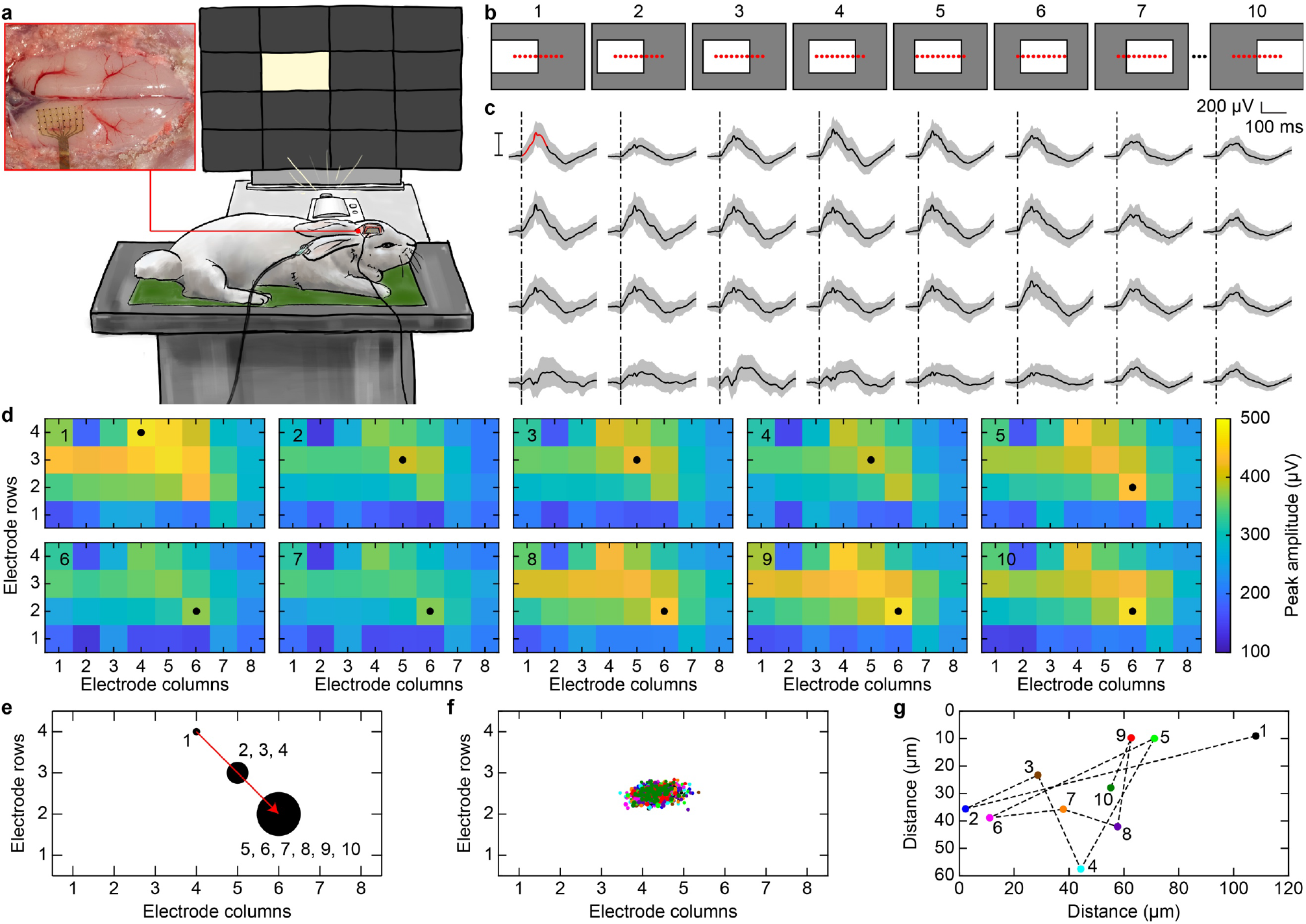
(**a**) Sketch of the experimental setup for visual stimulation. An anaesthetized rabbit is placed in front of the screen on which the light stimuli are projected. The red square shows an enlarged picture of the ECoG recording electrode array placed over V1 (post-mortem image). (**b**) Schematic representations of the patterned visual stimulation. The red dots are the centre of the white rectangles. (**c**) Representative example of the 32 VEPs recorded with the ECoG electrode array upon presentation of one visual stimulus. The black trace is the average of 60 repetitions, while the s.d. is the grey area. The trace in the top left corner shows the quantification of the peak-to-peak amplitude in the early portion of the signal (100 ms). The black dashed lines are the light stimulus onset. (**d**) Averaged activity maps among the 60 repetitions obtained for each visual stimulus location. In each map, the number in the upper left corner indicates the stimulus location, while the black circle indicates the leading channel. (**e**) Map of the channels identified as leading channels. The black circles’ size is proportional to the number of times the electrode is identified as the leading channel. The numbers next to the circles correspond to the stimulus locations leading to the electrode being identified as the leading channel. The red arrow identifies the leading channel’s shift. (**f**) Distributions of the centres of the activity map. Each circle corresponds to a single activity map upon one visual stimulus (60 circles for each stimulus location). Each colour corresponds to one stimulus location as in (**g**). (**g**) Shifts of the average centre of the activity maps.

Also, electrical stimulation elicits phosphenes, and a focalised flash is the closest stimulus to a phosphene. Despite these similarities, electrically evoked responses do not necessarily match visually evoked responses since electrical stimulation bypasses both phototransduction and the retinal network. Also, flashes are two orders of magnitude longer than electric pulses (16 ms and 150 μs, respectively).

Each circle is the average of the centres originating from a stimulus location. The numbers correspond to the stimulus location associated to a colour code.

One of the most commonly used approaches to determine what part of the visual cortex is activated by a stimulus is measuring the leading channel’s shift [46], which is the electrode with the highest peak-to-peak amplitude. This approach has been successfully used to determine retinotopic organisation in the visual cortex when paired with intracortical recordings. However, the low spatial resolution of ECoG microelectrodes might be inadequate for this task. In our dataset, we computed the activity map for each visual stimulus in each location as the peak-to-peak amplitude for each recording electrode. The 60 activity maps corresponding to a stimulus location have been averaged (**Figure 1d**), revealing that 3 leading channels appeared out of the 10 stimulus locations upon patterned visual stimulation (**Figure 1e**). The shift in the leading channel is coherent with the stimulus’s shift (from temporal to nasal) according to previously reported visual maps [50–52]. However, the limited spatial resolution of the recording system and analysis method provides only three hot-spots, which are not enough to identify distinguishable pattern of activation for each stimulus location. Therefore, we evaluated the whole activity maps. First, we computed the distribution of the centre of mass for each activity map (**Figure 1f**), and they resulted in overlapping distributions lacking significant clustering. Then, we computed the average centre of the distributions and evaluated the shift, which is in the order of one hundred microns and occurring in directions not coherent with the visual stimulus’s horizontal motion. These results indicate that the cortical activity map measured by the ECoG electrode array was not affected by the visual stimulus’s location.

The shifts of both the leading channel and the centre of mass alone are not a good estimator of the cortical activity pattern’s modulation upon variation of the stimulus location when recordings are made with ECoG microelectrodes, since they are both based only on the signal peak-to-peak amplitude. Therefore, we sought an alternative method encompassing more exhaustive information about the elicited cortical signal to identify the stimulation corresponding to a particular activation pattern in the visual cortex.

We investigated a machine-learning approach to isolate the most informative features leading to a finer analysis of the cortical signals, highlighting differences among signals generated by different stimuli. The hypothesis is that the similarity between signals will decrease when the areas of stimulation within the visual field are further apart, which will be reflected by higher classification accuracy. The machine-learning workflow classified the signal originating from patterned visual stimulation based on two types of features extracted from the 32 VEP time courses: continuous or punctual. The continuous features are the signal time course (‘Time’, **Figure 2a** in red), its derivative (‘Slope’, **Figure 2b** in red) and its broad-band gamma power (‘BGP’, **Figure 2c**). The punctual features, which have a single value per time course, are the peak-to-peak amplitude of the VEP signal (‘PA’) and the latency of the positive peak (‘PL’) (**Figure 2a**). In addition to the features derived from the time course, we included features containing information about the frequency domain to broaden the possible features. Therefore, we added continuous features consisting of each pair of time points and the power spectrogram frequency (which was down-sampled by a factor 10 in time). From the power spectrogram, we extracted three punctual features: amplitude (‘Power Amplitude’), frequency (‘Power Frequency’) and timing (‘Power Timing’) of the maximum of the power spectrogram (red cross in **Figure 2c**). Last, the wavelet power spectrogram was down-sampled in the time axis from 12 kHz down to 1200 Hz and used as the continuous feature (‘Power’). All the features were concatenated in a single matrix for feature selection.

**Figure 2.**
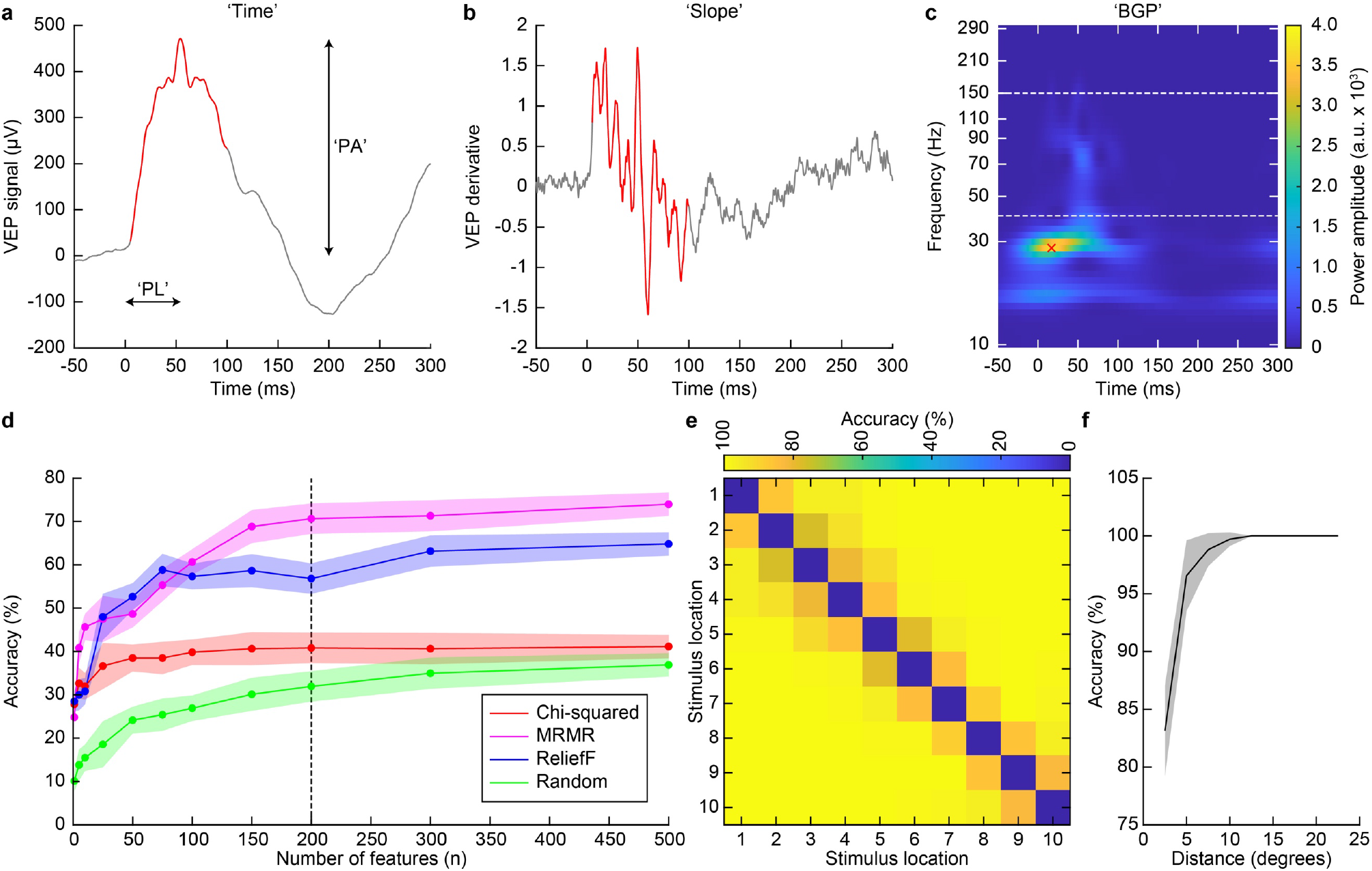
(**a**) Representative VEP signal highlighting the peak-to-peak amplitude and the positive peak latency. The red portion of the signal corresponds to the time course used as a feature from 5 ms to 100 ms. (**b**) Derivative of the VEP signal in **a**. The red portion of the signal is the one used as a feature from 5 ms to 100 ms. (**c**) Representative spectrogram from the VEP signal in **a**. The white dashed lines indicate the frequency band corresponding to the ‘BGP’. The red cross highlights the ‘Power Amplitude’, the ‘Power Frequency’ and the ‘Power Timing’. (**d**) Average (± s.d., 6 repetitions) classification accuracy of the feature selection algorithms. The black dashed line highlights the classification accuracy after 200 features. (**e**) Accuracy matrix for patterned visual stimulation. (**f**) Average (± s.d.) classification accuracy of the combination of pairs of stimuli as a function of the centre-to-centre distance between visual stimuli.

Since specific portions of the signal or some signal features are more informative than others, we applied a feature selection approach on the training set only. The very large number of features made some feature selection approaches computationally expensive, so we opted first to do a pre-selection test by computing the chi-score of all the individual features. We then proceeded to test different features selection algorithms (minimum redundancy maximum relevance, chi-squared test, reliefF and randomly selected features) on the 5% of the features with the highest chi-score (**Figure 2d**). We selected the minimum redundancy maximum relevance (MRMR) algorithm for our future analysis, as this algorithm resulted in the highest classification accuracy. When the number of selected features is disproportionally large compared to the number of observations, overfitting may occur: the RMSE of the training set converges to zero, while the RMSE of the testing set increases drastically. To avoid overfitting, we selected the best 200 features returned by the MRMR algorithm (one-third of the total number of observations) for which the RMSE of the testing set was minimized, and we used them as an input of a support machine vector classifier. The MRMR algorithm returned a classification accuracy 70.67% with 200 features (**Figure 2d**).

In a first step, we conducted the classification on all the stimuli at once (10 locations and 60 repetitions for a total of 600 observations), resulting in an average (± s.d.) classification accuracy of 71.17% ± 1.83% (10-classes classification). Then, to evaluate the impact of the distance (in degrees) between stimuli on the classification accuracy, we computed the classification accuracy between pairs of patterned visual stimuli (2-classes classification), resulting in 90 accuracy values organised into an accuracy matrix (**Figure 2e**). All the values separated by the same distance were averaged and plotted as a function of the distance (**Figure 2f**). We observed an increase in the classification accuracy when the distance between stimulus locations increased. The classification accuracy started from 83.15%, which is higher than the chance level (50%), monotonously increased for the lowest separations, reached 96% for a separation of 5 degrees, and 100% for separation higher than 7 degrees. Visual stimuli are 14 degrees wide, so 100% classification accuracy was reached when the stimuli’s overlap was reduced to 50%. This result suggested that the more the visual stimuli were separated, the less similar the VEPs are. In our hypothesis, as the patterned stimulations get further apart and gradually stop overlapping, they will recruit different populations of retinal ganglion cells, leading to the activation of increasingly different groups of neurons in V1. This activation ultimately results in a larger difference between the VEP signals, as they are an integrated summation of those neurons’ activity.

### Classification of electrical responses

We successfully designed a classification algorithm for patterned visual stimulation. Next, we tested it on the optic nerve’s intraneural electrical stimulation. First, we took advantage of a tailored OpticSELINE (Flat-OpticSELINE) with a linear and planar distribution of 8 electrodes transversally inserted from medial to lateral in the centre of the left optic nerve (**Figure 3a**). The advantage of this linear and flat electrode array is the similarity with the previously tested visual pattern: stimuli varying along a single dimension. Visual stimulation consisted of flashes aligned along the horizontal meridian from temporal to the nasal direction. With the Flat-OpticSELINE also electrical pulses are organised along the central line of the nerve’s cross-section. A liner electrodes’ distribution does not necessarily mean that phosphenes will occur along the horizontal meridian, since the fibres’ retinotopic distribution within the nerve is unknown. However, the liner electrodes’ distribution allowed us to compute the classification accuracy as a function of the distance and apply the linear regression model. Electrically evoked potentials (EEPs) were recorded upon biphasic charge-balanced asymmetric current pulses of different current amplitudes ranging from 10 to 2000 uA (**Figure 3b**).

**Figure 3.**
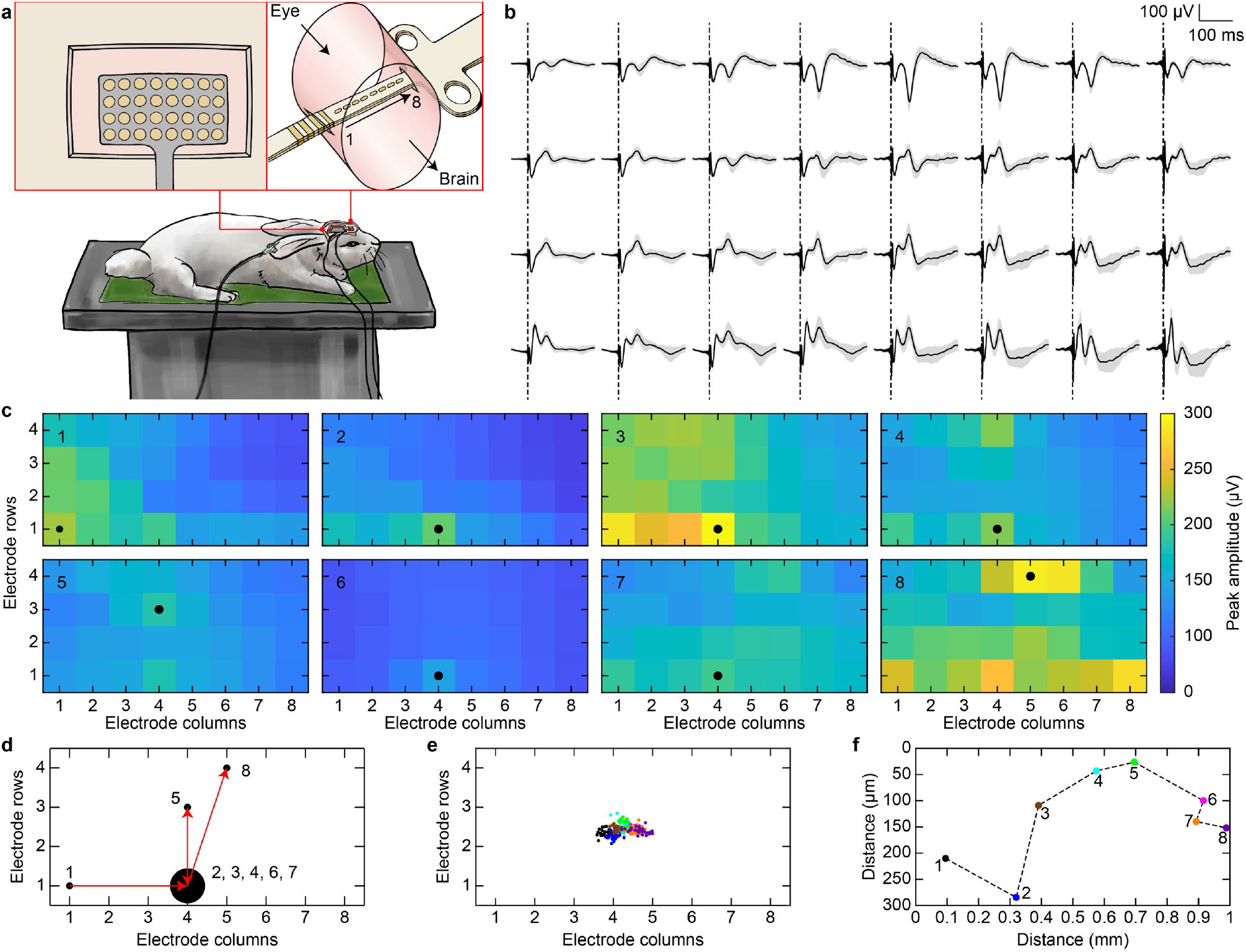
(**a**) Sketch of the experimental setup for electrical stimulation with the Flat-OpticSELINE. The red squares show an enlarged sketch of the ECoG recording electrode array placed over V1 and the Flat-OpticSELINE inserted into the left optic nerve from medial to lateral. The electrode’s arrangement is indicated by the black arrow from 1 to 8. (**b**) Representative example of the 32 EEPs recorded with the ECoG electrode array upon one electric pulse (2000 μA). The black trace is the average of 30 repetitions, while the s.d. is the grey area. The black dashed lines are the stimulus onset. (**c**) Averaged activity maps among the 30 repetitions obtained for each electrode upon electrical stimulation at 2000 μA. In each map, the number in the upper left corner indicates the electrode, while the black circle indicates the leading channel. (**d**) Map of the channels identified as leading channels upon electrical stimulation at 2000 μA. The black circles’ size is proportional to the number of times the electrode is identified as the leading channel. The numbers next to the circles correspond to the stimulation electrodes leading to the recording electrode being identified as the leading channel. The red arrows identify the shift of the leading channel. (**e**) Distributions of the centres of the activity map upon electrical stimulation at 2000 μA. Each circle corresponds to a single activity map upon one electric stimulus (30 circles for each electrode). Each colour corresponds to one stimulus location as in (**f**). (**f**) Shifts of the average centre of the activity map upon electrical stimulation at 2000 μA. Each circle is the average of the centres originating from one electrode. The numbers correspond to the electrode associated to a colour code.

First, we computed the activity maps for each stimulation electrode and each current amplitude (**Figure 4c**). From the maps, we computed the shift in both the leading channel (**Figure 3d**) and the average centre of mass (**Figure 4e**). At low current amplitudes (≤ 250 μA), both shifts appear random, but they became structured and reproducible from higher current amplitudes (≥ 750 μA). The quantification of the peak-to-peak amplitude from 750 μA revealed 2 to 5 leading channels out of the 8 stimulating electrodes depending on the current amplitude with a clear shift direction. Compared to the shift in VEPs, the direction is opposite, from posterior-lateral (electrode 1) to anterior-medial (electrode 8), which might be coherent with the insertion of the array from the medial to the lateral side of the nerve: electrode 1 is lateral, and electrode 8 is medial. Again, the limited spatial resolution of the recording system and analysis method provides only 2 to 5 hot-spots, which are not enough to identify distinguishable cortical activity patterns for each electrode. The distributions of the centre of mass for each activity map appeared more clustered and separated than the ones from VEPs (**Figure 3e**). Consequently, the shift of the average centre of the distributions from posterior-lateral (electrode 1) to anterior-medial (electrode 8) is larger than the one from VEPs. This larger shift and higher clustering could be justified since visual stimuli cover a small portion of the visual field, while electrodes spanned the entire optic nerve cross-section. We could hypothesise that electrodes induced phosphenes in region of the visual filed which are more distant than the visual stimuli.

**Figure 4.**
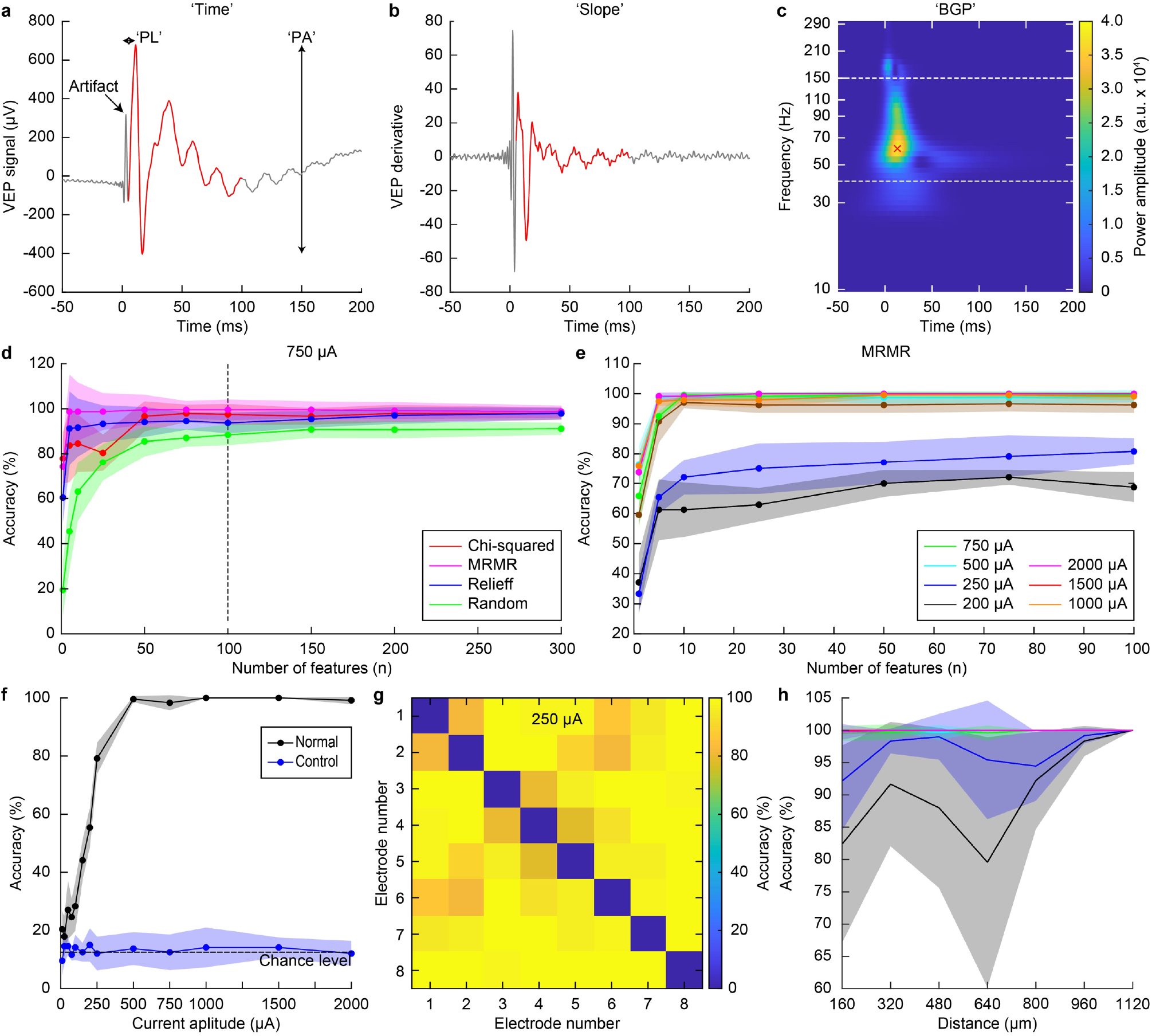
(**a**) Representative EEP signal highlighting the peak-to-peak amplitude and the latency of the positive peak. The red portion of the signal corresponds to the time course used as features from 5 ms to 100 ms. (**b**) Derivative of the EEP signal in **a**. The red portion of the signal is the one used as features from 5 ms to 100 ms. (**c**) Representative spectrogram from the EEP signal in **a**. The white dashed lines indicate the frequency band corresponding to the ‘BGP’. The red cross highlights the ‘Power Amplitude’, the ‘Power Frequency’ and the ‘Power Timing’. (**d**) Average (± s.d., 6 repetitions) classification accuracy of the feature selection algorithms for a stimulation current amplitude of 750 μA. The dotted line highlights the classification accuracy after 100 features. (**e**) Average (± s.d., 6 repetitions) classification accuracy of the MRMR feature selection algorithm for the stimulation current amplitudes from 200 μA to 2,000 μA. (**f**) Average (± s.d.) classification accuracy as a function of the stimulation current amplitude of all the electrodes (black). The blue circles and line show the classification accuracy when labels of the observations were scrambled. (**g**) Accuracy matrix for the optic nerve’s intraneural electrical stimulation at 250 μA. (**h**) Average (± s.d.) classification accuracy of the combination of electrodes’ pairs as a function of the centre-to-centre distance at increasing current amplitudes (from 200 to 2,000 μA) for the optic nerve’s intraneural electrical stimulation. Colour code is as in panel **e**.

For EEPs, we implemented the same classification workflow used for VEPs with the machine-learning algorithm. The features used and the feature selection method remained unchanged (**Figure 4a–c**), but due to the lower number of repetitions (30 instead of 60), we selected only the 100 best scoring features to avoid overfitting using the MRMR algorithm (**Figure 4d,e**). In a first step, we conducted the classification on all the stimuli at once (8-classes classification) for each current amplitude: 8 electrodes and 30 repetitions for a total of 240 observations (**Figure 4f**, black). The classification accuracy started around 20% for low current amplitudes (from 10 to 75 μA), slightly above the chance level (12.5% for 8 electrodes). This low correlation accuracy is justified since those current amplitudes correspond to the threshold to elicit a detectable EEP. The correlation accuracy linearly increased by increasing the current amplitude and reached nearly 100% from 500 μA. As a control experiment, the observations’ labels were randomly scrambled before the classification in both the training and testing sets, and the classification accuracy remained rightfully close to chance level regardless of the current amplitude (**Figure 4f**, blue). Overall, the machine-learning algorithm correctly classified cortical activity patterns based on the electrode in the array from which they were evoked in the entire range of current amplitudes above the activation threshold for EEP. Then, to evaluate the impact of the separation between electrodes on the classification accuracy of the elicited cortical activations, we computed the classification accuracy between pairs of electrodes (2-classes classification), resulting in 56 accuracy values organised into an accuracy matrix for each current amplitude (**Figure 4g**). All the values separated by the same distance were averaged and plotted as a function of the distance (**Figure 4h**). The classification accuracy with the machine-learning algorithm returned an increasing classification accuracy as a function of the electrode distance. The increase was marked for intermediate current amplitudes (200 and 250 μA), while at higher current amplitudes, the accuracy was quickly reaching 100%. For lower current amplitudes (below 150 μA), we could not find a clear relation between classification accuracy and electrode separation, probably because of the very low cortical activity elicited by such low currents.

The machine-learning algorithm returned an excellent classification accuracy for intraneural electrical stimulation with the Flat-OpticSELINE. Therefore, we generalised the approach and used the method to classify EEPs originating from the standard OpticSELINE having flaps distributing electrodes over the nerve’s cross-section. For this classification, we used a dataset obtained from previous experiments [47]. Briefly, the optic nerves of four rabbits were implanted with the OpticSELINE, and the six electrodes of the top shank were used for electrical stimulation. EEPs were collected with the same ECoG arrays (**Figure 5a**). For each of the four rabbits, we classified the time courses of the EEPs originating from the optic nerve’s stimulation using the six electrodes of the upper part of the OpticSELINE as a function of the current amplitude (**Figure 5b**). We conducted the classification on all the stimuli at once for each current amplitude: 6 electrodes and 10 repetitions for a total of 60 observations. The machine-learning algorithm showed a very high classification accuracy for three rabbits, close to 100% from 200 μA. In one rabbit (bottom left panel in **Figure 5b**), the classification accuracy was increasing with a slower slope, probably because higher currents were required to elicit detectable EEPs and the signal-to-noise ratio of the cortical responses was low. The average accuracy among the four rabbits started around 20% for low current amplitudes (10 and 25 μA), which is slightly above the chance level (16.6% for 6 electrodes). Then, the classification accuracy rapidly increased until reaching a plateau at 100% for the higher current amplitudes (**Figure 5c**).

**Figure 5.**
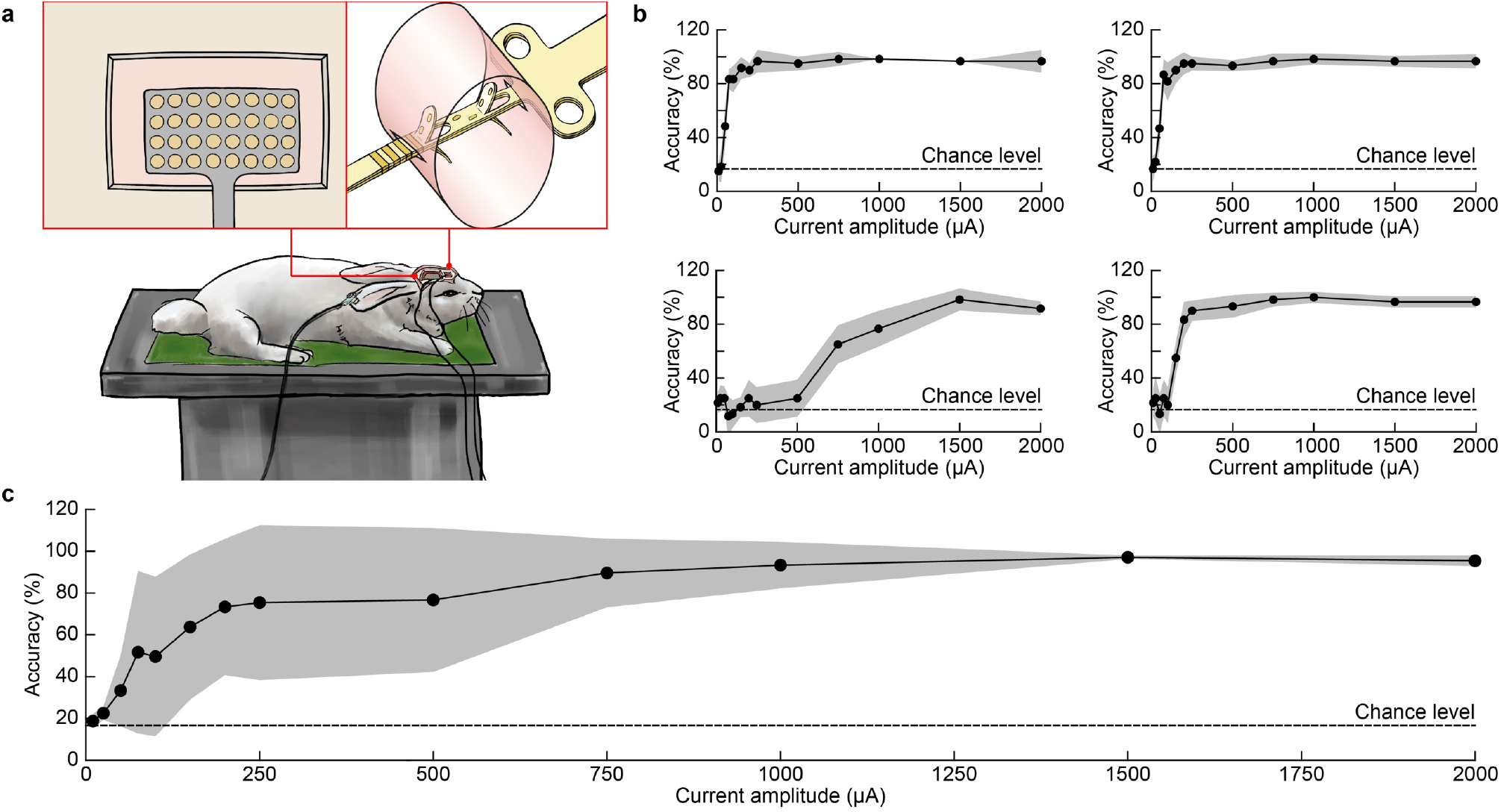
(**a**) Sketch of the experimental setup for electrical stimulation with the OpticSELINE. The red squares show an enlarged sketch of the ECoG recording electrode array placed over V1 and the OpticSELINE. (**b**) Average (± s.d.; 6 repetitions) classification accuracy as a function of the stimulating current for each rabbit. (**c**) Mean (± s.d.; *n* = 4 rabbits) classification accuracy as a function of the current amplitude.

### Linear regression of patterned visual and electrical stimulation

While the classification algorithm could classify cortical patterns originating from visual and electrical stimuli in different locations with high accuracy, the classifier had one limitation. If giving as input to the classifier a novel signal from an unknowing origin, it would only associate it with one of the classes of signals already encountered in its training phase, but it would not extrapolate the actual location of the stimulus eliciting that signal. This limitation motivated us to build a linear regression model to improve the predictive capacity when presented with a new signal of unknown origin, using the features with the highest individual R^2^ scores: 200 for patterned visual stimulation and 100 for patterned electrical stimulation.

For patterned visual stimulation, the prediction of the model was close to the actual centre of the rectangles (**Figure 6a**), with an average (± s.d.) RMSE of 2.72 ± 0.60 degrees (**Figure 6d**), equivalent to 12.45% of the full range of possible positions. The model performed better for the central locations, while the performance slightly decreased at the extremities. The adjusted R^2^ score of 0.86 suggested that the linear model is a good fit for patterned visual stimulation. To validate the model further, we performed two control experiments. First, we randomly scrambled the labels of the stimulus’s centres in both the training and testing sets (scrambled condition), where all stimuli corresponding to one location have been randomly assigned to another location (e.g. location 1 at −11 degrees was renamed +11 degrees). The linear regression model had a worst performance to predict the positions of the stimuli (**Figure 3b**), with an average (± s.d.) RMSE of 6.29 ± 2.16 degrees (**Figure 3e**) and very large variation in the average difference. Second, we removed one location from the training set, and we trained on the 9 remaining locations. The testing set was composed only by the location that was left out from the training set (left-out condition). The linear regression model correctly predicted the missing stimulus position (**Figure 3c**), with an average (± s.d.) RMSE of 2.90 ± 0.96 degrees (**Figure 3f**).

**Figure 6.**
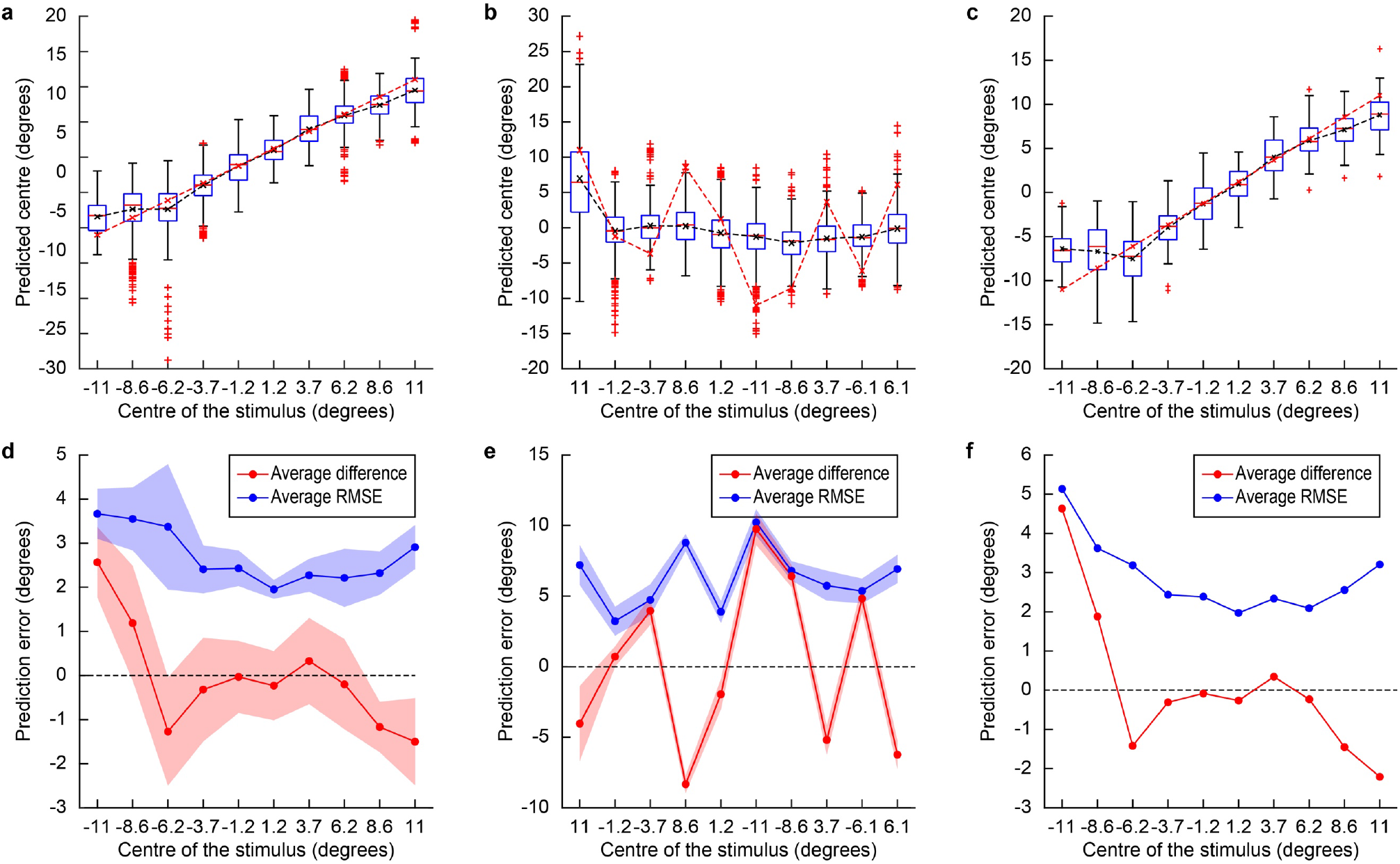
(**a,b,c**) Quantification of the averaged predicted centre locations (black dashed line) and their actual locations (red dashed line) for the three conditions: normal (**a**), scrambled (**b**) and left-out (**c**). In each box: the red line is the median, the box extremities are the 1^st^ (q1) and 3^rd^ (q3) quartiles, the whiskers are q1 + 1.5 × (q3 – q1) and q3 + 1.5 × (q3 – q1) and the red crosses outside of the whiskers are the outliers. (**d,e,f**) Average difference between the predicted and the actual locations (red) and RMSE of the predicted locations (blue) for the three conditions: normal (**d**), scrambled (**e**) and left-out (**f**). In panels (**d,e**) the shaded area is the s.d. In the left-out condition (**f**), the RMSE is computed only once for each excluded stimulus, so the s.d. cannot be computed.

Next, we fitted the linear regression model on the features extracted from the EEPs (**Figure 7**). The model had the best fit for 1,000 uA (**Figure 7a,b**) according to the adjusted R^2^ (0.91) and the average RMSE (0.68). Overall, among the various current amplitudes, the model had an adjusted R^2^ between 0.01 and 0.91, increasing with the current amplitude and an average RMSE between 3.24 and 0.65, decreasing with the current amplitude (**Figure 7c**). In the scrambled condition, the linear regression was worst (**Figure 7d**), with an average (± s.d.) RMSE of 2.06 ± 0.53 (**Figure 7e**). However, it can be noted that while some electrodes are not accurately predicted, other they still are despite the random scrambling. A similar behaviour was observed at all current amplitudes (**Figure 7f**). In the left-out condition, the linear regression model correctly predicted the missing electrode position (**Figure 7g**), with an average (± s.d.) RMSE of 1.06 ± 0.74 (**Figure 7h**). Overall, among the various current amplitudes, the model had an average RMSE between 4.06 and 1.06, decreasing with the current amplitude (**Figure 7i**).

**Figure 7.**
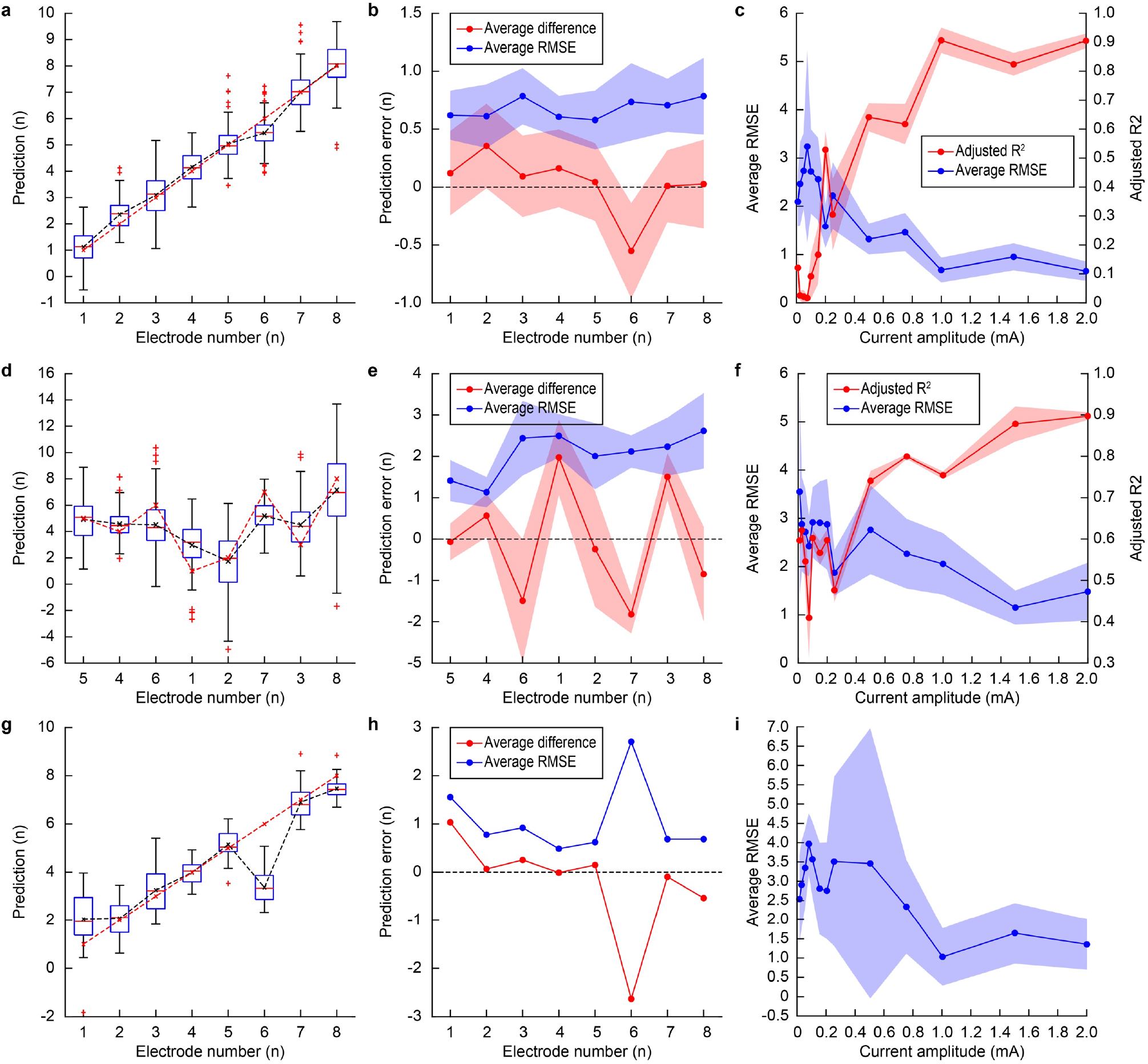
(**a,d,g**) Quantification of the averaged predicted centre locations (black dashed line) and their actual locations (red dashed line) upon electrical stimulation at 1,000 μA for the three conditions: normal (**a**), scrambled (**d**) and left-out (**g**). In each box: the red line is the median, the box extremities are the 1st (q1) and 3rd (q3) quartiles, the whiskers are q1 + 1.5 × (q3 – q1) and q3 + 1.5 × (q3 – q1) and the red crosses outside of the whiskers are the outliers. (**b,e,h**) Average difference between the predicted and the actual locations (red) and RMSE of the predicted locations (blue) upon electrical stimulation at 1,000 μA for the three conditions: normal (**b**), scrambled (**e**) and left-out (**h**). In panels (**b,e**) the shaded area is the s.d. In the left-out condition (**h**), the RMSE is computed only once for each excluded stimulus, so the standard deviation cannot be computed. (**c,f,i**) Average (± s.d.) RMSE (blue) and adjusted R^2^ coefficient (red) of the model as a function of the stimulating current amplitudes for the three conditions: normal (**c**), scrambled (**f**) and left-out (**i**). In the left-out condition (**i**), the adjusted R^2^ coefficient cannot be computed since it requires the variance of the testing set labels, which is null since the testing set is only composed of observations with the same label.

Our approach used nine classes of features from both the time domain and the frequency domain classified in two major categories: punctual and continuous. Therefore, we investigated which feature’s types was most informative. We repeated the linear regression model for both patterned visual stimulation and patterned electrical stimulation. We computed the correlation coefficient between the features and the stimuli’ locations based on the whole datasets to be more descriptive (i.e. on the total number of observations instead of the 5/6th of the training sets’ observations). Then, we selected the 200 (for patterned visual stimulation) and 100 (for patterned electrical stimulation) features with the highest correlation coefficient value (**Figure 8**). For patterned visual stimulation, the 200 selected features were from only two types: ‘Slope’ (51.5%) and ‘Power’ (48.5%), both continuous features. This result indicates that continuous features might be more informative than punctual features, like the peak-to-peak amplitude. Patterned electrical stimulation revealed a remarkably similar behaviour. Among all the current amplitudes, 6 types of features were cumulatively selected: ‘Time’ (8.5%), ‘Slope’ (12.5%), ‘Power Timing’ (0.2%), ‘Power Amplitude’ (0.2%), ‘BGP’ (19.2%) and ‘Power’ (59.4%). Continuous features (99.6%) were largely predominant compared to punctual features (0.4%). Features in the frequency domain were mostly selected (79%) compared to features in the time domain (21%). The most informative feature was the ‘Power’ feature (59.4%).

**Figure 8.**
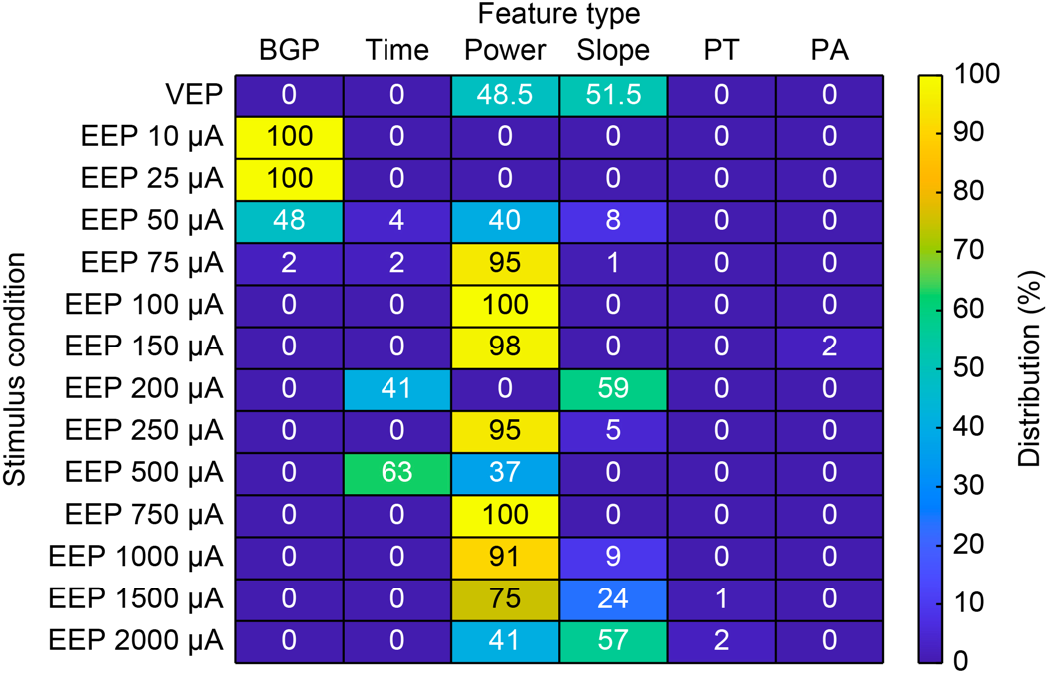
Distribution of the selected feature’s types. Numbers in each box correspond to the exact percentage value. PT corresponds to ‘Power Timing’ and PA to ‘Power Amplitude’.

## Discussion

Optic nerve stimulation has advantages compared to epiretinal stimulation. However, one of the key open questions in preclinical studies is to detect distinguishable cortical patterns upon selective activation of the nerve fibres. Computational and experimental methods showed that intraneural electrodes could selectively activate different sets of fibres within the optic nerve [47]. Also, high-frequency pulse trains with a variable number of pulses could modulate the cortical responsivity without affecting the optic nerve’s activated area. Building on this evidence, our results showed that a machine-learning algorithm could correctly classify electrically evoked cortical potentials originating from the microelectrodes of the OpticSELINE array.

First, we showed that approaches relying only on the peak-to-peak amplitude alone are not powerful enough to distinguish the stimulus’s location inducing the cortical activation due to the low spatial resolution of ECoG microelectrodes. This result motivated the development of a new signal analysis approach based on a SVM classification algorithm, and we took advantage of patterned visual stimulation for its validation. We designed the stimulation pattern to test conditions in which the visual field’s stimulated portions are close and overlapping, thus more difficult to distinguish from one another. We hypothesised that the further apart the visual field’s stimulated portion would be, the less similar the signal originating from their activation would be. Therefore, the visual stimuli were aligned over the horizontal meridian to precisely quantify their centre-to-centre separation. The SVM classification method confirmed the hypothesis, showing an increase in classification accuracy with increased centre-to-centre separation, reaching 100% accuracy for stimuli with less than 50% overlap.

After demonstrating the efficiency of the SVM classification approach on patterned visual stimulation, we tested it on the optic nerve’s intraneural electrical stimulation. First, we used a simplified intraneural electrode array (Flat-OpticSELINE). As for visual stimulation, we found an increase in classification accuracy with increased centre-to-centre separation between the stimulating electrodes. However, the performance of the SVM classification method was much better for electrical stimulation than visual stimulation, with nearly 100% classification accuracy from the smallest separation distance above 500 μA. The better performance on electrical stimulation can be explained since the visual stimuli cover a small portion of the visual field, while electrodes spanned the entire optic nerve. We could hypothesise that electrodes induced phosphenes in region of the visual filed which are more distant than the visual stimuli. Because of this difference, a quantitative comparison between visual and electrical stimulation could not be performed. However, both modalities showed the same qualitative behaviour. This result also highlighted the classification accuracy’s dependence on the current amplitude, with higher accuracy at higher amplitudes. Higher current amplitudes activate a larger portion of the nerve, increasing the overlap between neighbouring electrodes. We hypothesised that the classification accuracy still increases because the nerve’s activated portion remains sufficiently different, leading to distinguishable cortical patterns. Also, the higher is the current amplitude, the higher is the signal-to-noise ratio of the recorded signal, and the lower is the variability among trials, leading to better classification.

To generalise the approach, we implemented the SVM classification algorithm to EEPs induced by the OpticSELINE intraneural array, where the electrodes are not distributed in line but cover the cross-section of the nerve. The algorithm returned similar results, with a nearly 100% classification accuracy above 200 μA of current stimulation in three out of the four rabbits tested. According to the Shannon criteria [53], the OpticSELINE can safely deliver current pulses up to 500 μA (k = 1.85). In one animal, the classification accuracy increased with a slower slope as a function of the current. This result was explained since, in that experiment, higher currents were required to elicit detectable EEPs, and the signal-to-noise ratio of the cortical responses was low. The classification accuracy’s dependency with the signal-to-noise ratio makes the classification algorithm relative to the stimulation strength. Thus, an increase in classification accuracy could be linked to a larger separation between the stimuli only when the stimuli are comparable in strength: that is to say, the same illuminated surface for visual stimuli or the same current amplitude for electrical stimulation.

The SVM algorithm provided only a qualitative estimation of the distance separating two stimuli: a low accuracy score was associated with a small distance between the stimuli, while a high accuracy score indicated that the stimuli were further apart. However, it cannot predict how far apart the stimuli were since after a small distance between the electrodes, the classification accuracy reached a plateau at 100% of accuracy, and increasing the distance between stimuli further could note improve it. Also, for electrical stimulation with the Flat-OpticSELINE above 500 μA, the classification accuracy reached 100% from the smallest separation distance. Therefore, we applied a regression model to add a predictive capacity. We selected a linear model as it is a simple approach matching the type of stimuli used, which vary along a single dimension. This choice relies on the assumption that some features are linearly related to the stimuli’ coordinates. We obtained several features with a large correlation coefficient, which allowed us to build a successful regression model for both visual and electrical patterned stimulation. The linear regression model performed better for patterned visual stimulation than patterned electrical stimulation, because in visual stimulation the stimuli remained located in a small and central portion of the visual field. The approximation of linearity is less accurate for patterned electrical stimulation since electrodes are located through the whole size of the optic nerve, and the evoked phosphenes might not be all linearly organised. In this condition, a more complex machine-learning algorithm such as convolutional neural networks could provide better results since it would not impose any assumption on linearity or any other type of regression.

The linear regression models for both patterned visual and electrical stimulation revelated that continuous features are the most informative. Punctual features appeared only sporadically in pattered electrical stimulation and resulted in not being very informative. Punctual features compress the cortical signal into a single value, which does not appear to be informative enough for prediction models. Features from both time and frequency domains were selected. However, spectral features appeared to be the most robust, especially the ‘Power’ feature. Among features in the time domain, ‘Slope’ was the most selected. It is noteworthy that the signal’s derivative (‘Slope’) was mostly selected and not the signal itself (‘Time’). This result might be explained by the trial-to-trial variation of the signal (e.g. voltage offset), which is removed in the derivative, making the ‘Slope’ more consistent between repetitions. Among features in the frequency domain, it appears that the ‘Power’ feature is the most informative. BGP or similar frequency ranges have been used to successfully classify cortical signals in V1 upon visual stimulation [49] and electrical stimulation of the retina [54]. From this dataset and results, the whole spectrum ('Power’ feature) is more informative than its partition (‘BGP’). The BGP might results in a loss of information compared to the whole spectrum.

The successful design of these two algorithms shows that the cortical activities elicited by the electrical stimulation are indeed distinctively different. The linear regression model then shows that they are meaningfully different, as it highlights features that vary in a linear manner, which can be expected from cortical activity patterns resulting from the stimulation of a gradually shifted portion of the visual field.

## Conclusion

The machine-learning approach based on a SVM classification algorithm showed that the cortical activity patterns induced by the optic nerve’s intraneural stimulation with the OpticSELINE could be successfully classified. Like patterned visual stimulation, intraneural electrodes might selectively recruit different optic nerve portions leading to different visual phosphenes. This result is a critical step to move forward visual prosthesis based on the optic nerve’s intraneural stimulation. However, it must be noted that this analysis and results cannot provide information about perceptual phenomena. Also, in late-stage retinitis pigmentosa patients, only 30% of retinal ganglion cells might be spared, and the retina’s hyperexcitability might affect the reliability of artificial vision via optic nerve’s stimulation. These assessments will require further clinical investigation.

## Acknowledgement

This work was supported by École polytechnique fédérale de Lausanne, Medtronic and Swiss National Science Foundation (200021_182670).

V.G performed data analysis. E.B. fabricated the devices and performed experiments. E.G.Z. performed experiments. D.G. designed and led the study, and wrote the manuscript. All the authors read and accepted the manuscript.

The authors declare no competing interests.

